# Cullin-5 regulates nuclear positioning and reveals insights on the sensing of the nuclear-to-cytoplasmic ratio in *Drosophila* embryogenesis

**DOI:** 10.1101/2022.01.24.477546

**Authors:** Luke Hayden, Anna Chao, Victoria E. Deneke, Alberto Puliafito, Stefano Di Talia

## Abstract

In most metazoans, early embryonic development is characterized by rapid division cycles which pause before gastrulation at the mid-blastula transition (MBT).^1^ These early cleavage divisions are accompanied by cytoskeletal rearrangements which ensure proper nuclear positioning. Yet, the molecular mechanisms controlling nuclear positioning are not fully elucidated. In *Drosophila*, early embryogenesis unfolds in a multinucleated syncytium, and nuclei rapidly move across the anterior-posterior (AP) axis at cell cycles 4-6 in a process driven by actomyosin contractility and cytoplasmic flows.^2,3^ Previously, *shackleton* (*shkl*) mutants were identified in which this axial spreading is impaired.^4^ Here, we show that *shkl* mutants carry mutations in the *cullin-5* (*cul-5*) gene. Live imaging experiments show that Cul-5 is downstream of the cell cycle but required for cortical actomyosin contractility. The nuclear spreading phenotype of *cul-5* mutants can be rescued by reducing Src activity genetically, suggesting that a major target of Cul-5 is Src kinase. *cul-5* mutants display gradients of nuclear density across the AP axis at the MBT which we exploit to study cell cycle control as a function of the N/C ratio. We found that the N/C ratio is sensed collectively in neighborhoods of about 100μm and such collective sensing is required for a precise MBT in which all the nuclei in the embryo pause their division cycle. Moreover, we found that the response to the N/C ratio is slightly graded along the AP axis. These two features can be linked to the spatiotemporal regulation of Cdk1 activity. Collectively, our results reveal a new pathway controlling nuclear spreading and provide a quantitative dissection of how nuclear cycles respond to the N/C ratio.

## Results

### *shkl* encodes the ubiquitin ligase *cullin-5*

*shkl* mutants are among the few genetic perturbations which have been shown to directly impinge on the spreading of nuclei in early *Drosophila* embryogenesis.^4,5^ Moreover, *shkl* embryos display gradients in nuclear density, which have been linked to a significant decrease in the synchrony of the last cell cycle preceding the MBT.^3^ To elucidate how *shkl* regulates nuclear positioning and how such regulation impacts cell cycle lengthening at the MBT, we first obtained two *shkl* alleles identified in the original mutagenesis screen (*shkl*^*GM130*^ and *shkl*^*GM163*^) and imaged embryos laid by transheterozygous mothers (hereinafter *shkl* embryos). We confirmed that in *shkl* embryos the nuclear cloud failed to reach the posterior pole of the embryo at the correct time and nuclei were not positioned uniformly, as seen previously (Fig. 1A-B).^4^ Moreover, we found that the lower density of nuclei in the posterior of the embryo was frequently accompanied by an extra round of nuclear divisions (Supplementary Movie 1). Thus, failures in nuclear positioning can have significant impact on the collective and synchronous decision of all nuclei to remodel the cell cycle at the MBT.

**Fig. 1.**
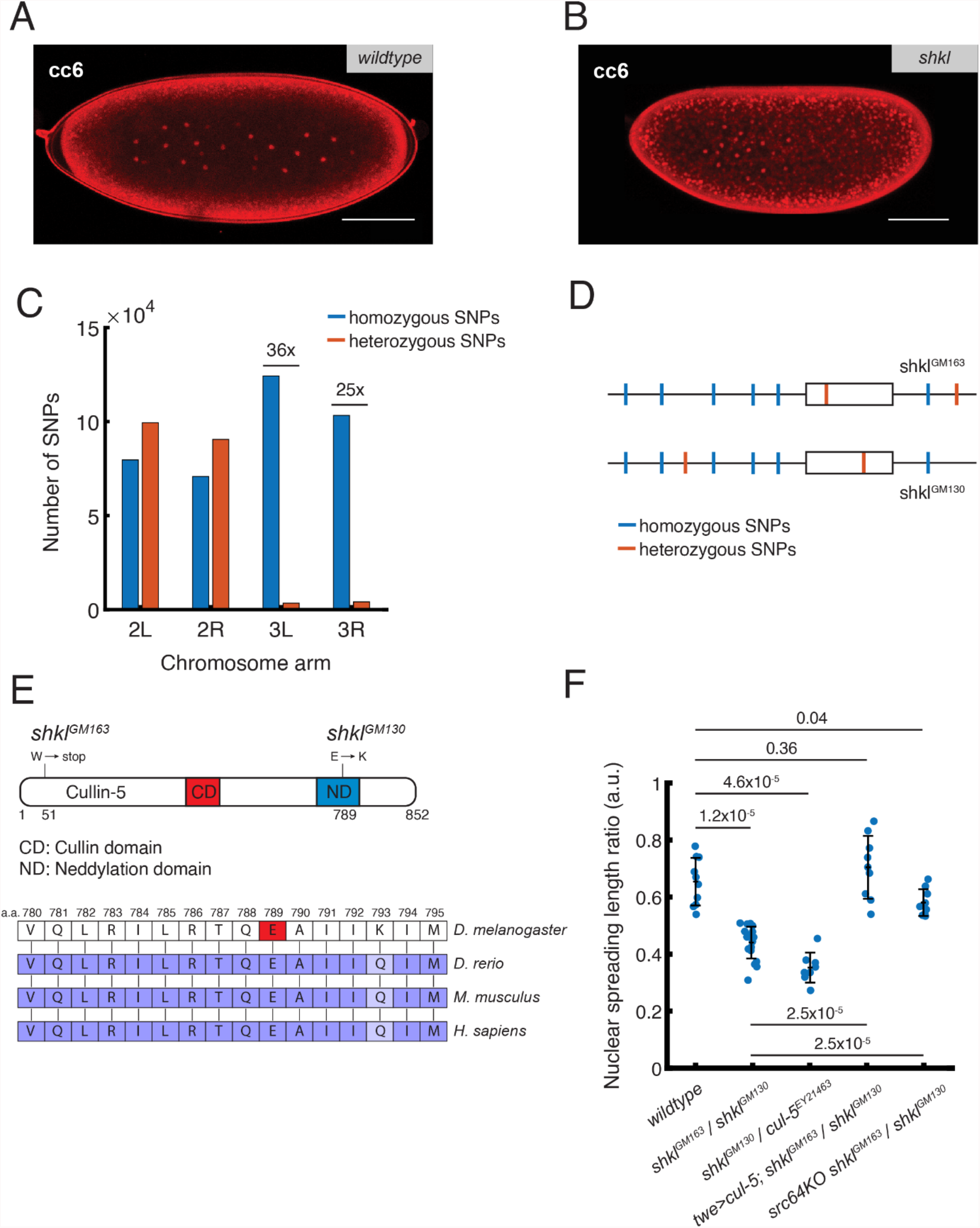
Genetic identification of *shkl* mutants. (A, B) Nuclear positioning in interphase of cc 6 in wildtype (A) and *shkl* (B) embryos, visualized with PCNA-TagRFP. (C) Number of homozygous and heterozygous SNPs (relative to the reference genome) in *shkl* embryos in each arm of chromosome two and three. The mutant screen^4^ was performed on an isogenic third chromosome and thus only the third chromosome has reduced heterozygous SNPs. (D) A theoretical example gene with heterozygous SNPs between the two *shkl* alleles. (E) Top: Schematic for the Cul-5 protein with domains and *shkl* mutations shown. The *GM130* allele of *shkl* contains a point mutation in the neddylation domain where it is evolutionarily conserved. The *GM163* allele contains a point mutation leading to an early stop codon. Bottom: Part of the neddylation domain of the Cul-5 protein in different model organisms. Highlighted residue indicates *GM130* mutation site. (F) Genetic complementation and rescue tests. Fixed embryos were stained with DAPI to show nuclear positioning at cc 6. The ratio between the length of the nuclear cloud and the length of the embryo was measured. Data are represented as mean ± SEM. P-values (Kolmogorov–Smirnov test) are shown. Each dot represents one embryo. *cul-5*^*EY21463*^ is a hypomorphic mutant allele. twe>cul-5 is a transgenic line carrying a plasmid with cul-5 cDNA under the regulation of the twine promotor. Scale bar: 100μm.

To identify the *shkl* gene, we used a DNA sequencing approach, centered on the fact that the original screen was performed in a strain carrying an isogenic third chromosome and that two alleles of the *shkl* gene were available.^4^ We reasoned that the third chromosomes of these two alleles had little time (∼20 years) to accumulate mutations with respect to each other. Thus, we predicted that genomic sequencing of *shkl* flies (*shkl*^*GM130*^/*shkl*^*GM163*^) would show a much lower number of heterozygous single nucleotide polymorphisms (SNPs) than homozygous ones relative to the reference genome on the third chromosome, a prediction which we could readily confirm (Fig. 1C). Taking advantage of the low number of heterozygous SNPs and the previous mapping^4^ of *shkl* between two markers (*ebony* and *claret*) on the right arm of chromosome 3, we looked for genes that carried two heterozygous missense SNPs with the idea that this would narrow our search to only a handful of genes (Fig. 1D). Bioinformatics analysis confirmed the validity of this argument, and we identified *cullin-5* (Cul-5) as the best candidate as allele *shkl*^*GM163*^ carried a premature stop codon at amino acid 51 and *shkl*^*GM130*^, a missense mutation (E to K) in the very conserved neddylation domain, a domain required for ubiquitin ligase activity and which is well-conserved across evolution (Fig. 1E).^6^ To perform complementation and rescue experiments, we fixed and stained embryos with DAPI to estimate the extent of nuclear spreading by measuring the shape of the nuclear cloud in cell cycle (cc) 6. We found that *shkl* alleles failed to complement an available *cullin-5* mutant (Fig. 1F). In addition, maternal expression of *cullin-5* from the *twine* promoter (which drives expression specifically in the germline^7^) was able to significantly rescue the nuclear spreading defects (Fig. 1F). Collectively, these results identify *shkl* mutants as alleles of the *cullin-5* genes and demonstrate that maternal expression of *cullin-5* is important for nuclear positioning in *Drosophila* embryos.

### *shkl* is downstream of the cell cycle and regulates cortical contractility

In our previous work, we showed that nuclear spreading is driven by cytoplasmic flows generated by cortical actomyosin contractility, which is in turn controlled spatiotemporally by the cell cycle oscillator (Fig. 2A).^3^ To quantify the degree to which cytoplasmic flows are disrupted in *shkl* embryos, we used yolk autofluorescence images to perform particle image velocimetry (PIV) in live embryos also expressing PCNA-TagRFP (which allows visualization of nuclei deep in the embryo) and measured the velocity of the cytosol and nuclei during the early cell cycles when axial expansion occurs. As previously shown, the wildtype embryos showed strong cytoplasmic flows coupled with nuclear movement which spread the nuclei across the anterior-posterior (AP) axis by the end of cell cycle 6 (Fig. 2B, D, S1).^3^ In contrast, cytoplasmic flows and nuclear movement in *shkl* embryos were sharply reduced (Fig. 2C, E, S1). Since cytoplasmic flows are generated by recruitment of active Myosin II to the cortex by active Rho,^3,8^ we sought to determine if the activities of these regulators are perturbed in *shkl* embryos. To that end, we measured the dynamics of a Rho biosensor^9^ and myosin II recruitment to the embryo cortex. Both Rho activity (Fig. 2F, S1) and myosin II recruitment (Fig. 2G) were reduced in *shkl* embryos as compared to wildtype. Next, we analyzed whether Cul-5 might impact actomyosin contractility by regulating the cell cycle. To this end, we looked at the master regulator Cdk1 by measuring the Cdk1-to-PP1 activity ratio using a FRET-based biosensor in both wildtype and *shkl* embryos.^10,11^ The oscillations in the activity ratio were similar in wildtype and *shkl* embryos. Moreover, the duration of the cell cycle near the middle of the embryo was also essentially unaltered (Fig. 2H, S2). Therefore, we argue that Cul-5 does not regulate the cell cycle oscillator. Taken together, our results indicate that Cul-5 is necessary for the proper activity of Rho and recruitment of myosin II which in turn regulate cortical contractility and nuclear positioning.

**Fig. 2.**
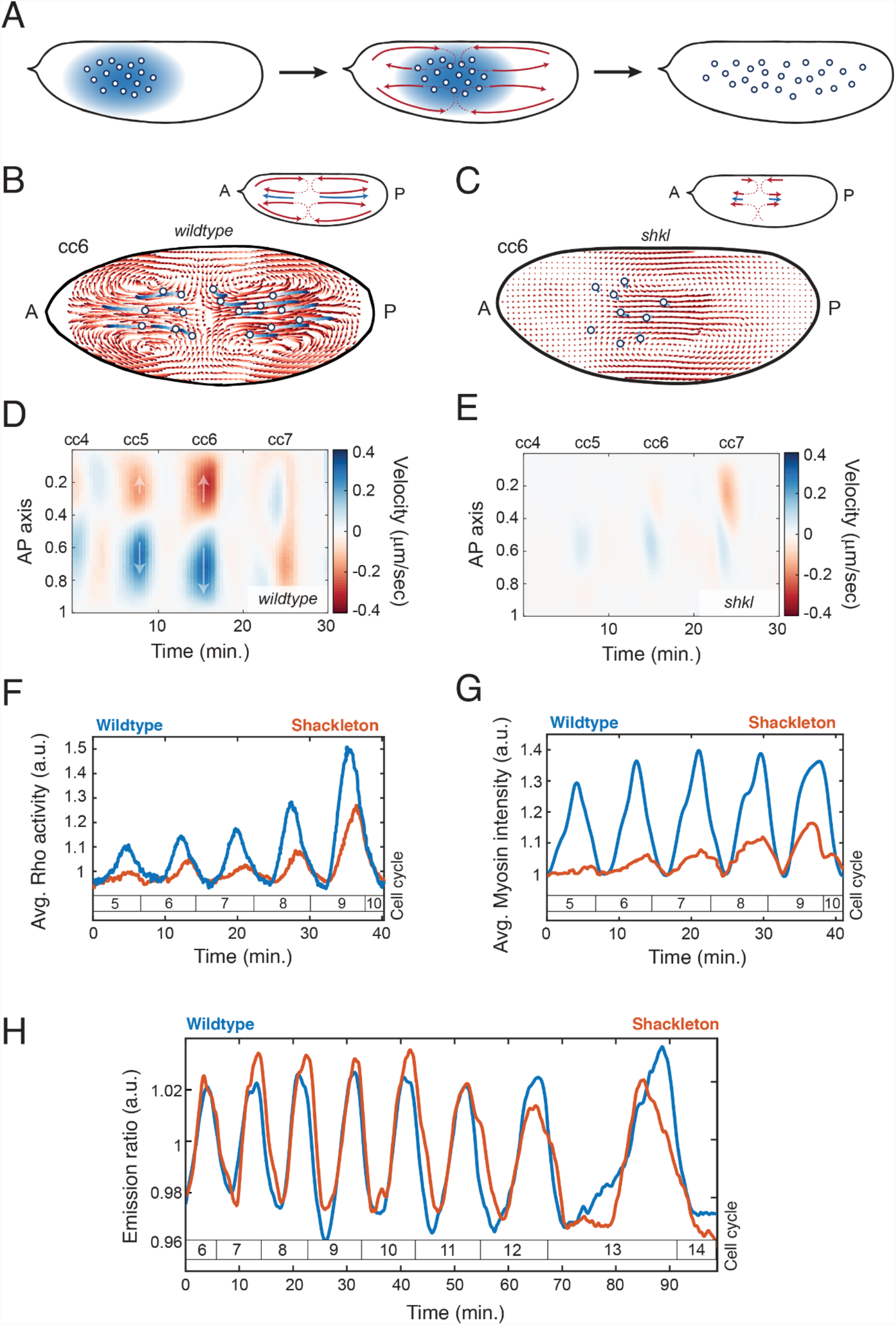
Characterization of the Cul-5 pathway. (A) Pathway for self-organized nuclear positioning in wildtype embryos; PP1 activity spreads from nuclei to the embryo cortex where it leads to gradients of myosin accumulation, thus generating cytoplasmic flows. These flows position nuclei uniformly across the AP axis. (B, C) PIV in wildtype (B) and *shkl* (C) embryos showing reduced cytoplasmic flows and nuclear movement in *shkl* embryos. Top insets depict magnitude and directionality of flows. (D, E) Heatmap quantification of the cytoplasmic flows in wildtype (D) and *shkl* (E) embryos. Arrows depict directionality of flows. (F, G) Both cortical Rho activity (F) and cortical myosin accumulation (G) are reduced in *shkl* embryos. (H) Cortical oscillations of the Cdk1/PP1 ratio from the FRET biosensor are similar in wildtype and *shkl* embryos.

### Cul-5 regulates cortical contractility through restricting the activity of Src

Cul-5 is a ubiquitin ligase which works in conjunction with other factors to regulate protein stability.^12^ A major target of Cul-5 is Src kinase^13-15^, whose activity is restricted by several ubiquitin ligases.^16-18^ Src is known to regulate the cytoskeleton^19,20^, including actomyosin contractility.^21^ These observations suggest that Cul-5 could (at least partly) regulate the axial expansion process by restricting Src activity. To test this hypothesis, we performed experiments using the Gal4/UAS system to overexpress constitutively active forms of the two Src homologs in *Drosophila*, Src42A or Src64B.^22^ We saw that overexpression of either homolog is sufficient to recapitulate the *shkl* phenotype with sharply reduced cytoplasmic flows and nuclear spreading (Fig S1). Similarly, if *shkl* embryos have reduced cortical contractions due to excessive Src activity, then genetically decreasing Src activity should rescue the *shkl* phenotype. We examined *shkl* mutants which also had only one copy of Src64B (heterozygous for *Src64B*^*KO*^, described previously^20^) and quantified cytoplasmic flows and nuclear positioning. We saw that reducing Src activity in *shkl* embryos significantly reduced the defects in axial expansion (Fig. 1F). Collectively, these results implicate the Cul-5/Src cascade in the regulation of nuclear positioning in *Drosophila* embryos.

### Nuclei sense the local nuclear density in large groups to determine whether to divide

In wildtype embryos, the morphogenetic processes driving nuclear positioning ensure that nuclear density is rather uniform across the embryo.^3^ Since nuclear divisions are synchronized within minutes^10,23^ by Cdk1 waves,^10,24^ nuclear density increases in 2-fold increments, which likely facilitates the consistent cell cycle lengthening observed in all nuclei at the MBT. To gain insights on how the embryo can achieve a robust response to changes in the N/C ratio, we observed that the nuclear spreading defects in *shkl* embryos cause a nuclear density gradient across the AP axis with lower density at the posterior (Fig. 3A-B, Fig. 4A). Therefore, we exploited this gradient to probe the response to gradual changes in nuclear density. We observed that the lower density of nuclei in the posterior of the embryo is frequently accompanied by an extra round of nuclear division (Fig. 3C). Interestingly, the size of the region which does an extra division varies across embryos and is frequently less than half the size of the embryo. Moreover, a salt-and-pepper phenotype with many regions of extra divisions next to regions of normal divisions is never observed.^25,26^ These two observations suggest that the nuclei do not sense the N/C ratio globally or in an autonomous manner, but rather they do so in a collective manner over some distance, here called the community radius. To infer this radius, we divided embryos into grids, measured the nuclear density within circles of different radii, and scored whether each grid point was within a region of either normal or extra division (Fig. 3D). We then fit these curves to logistic equations and used the N/C ratio at which the curves crossed 50% probability as the best N/C ratio threshold predictor, above which we predict embryos do not divide (Fig. 3E). As previously reported, these thresholds were around 70% of the wildtype cell cycle 14 nuclear densities.^25^ This value is half-way between the nuclear density at cell cycle 13 and 14, which likely contributes to the decision of all nuclei to divide rapidly at cell cycle 13 and lengthen their cycle 14. Next, we asked what is the fraction of nuclei that would fail to lengthen their cell cycle at the MBT as a function of the community radius. We found that to ensure that all nuclei (>99%) undergo a collective pause at the MBT, the response to the N/C ratio must be averaged over a community radius of at least 35μm (Fig. 3F). Such a community would contain about one hundred nuclei, thus implying that a robust cell cycle decision at the MBT requires a collective nuclear response.

**Fig. 3.**
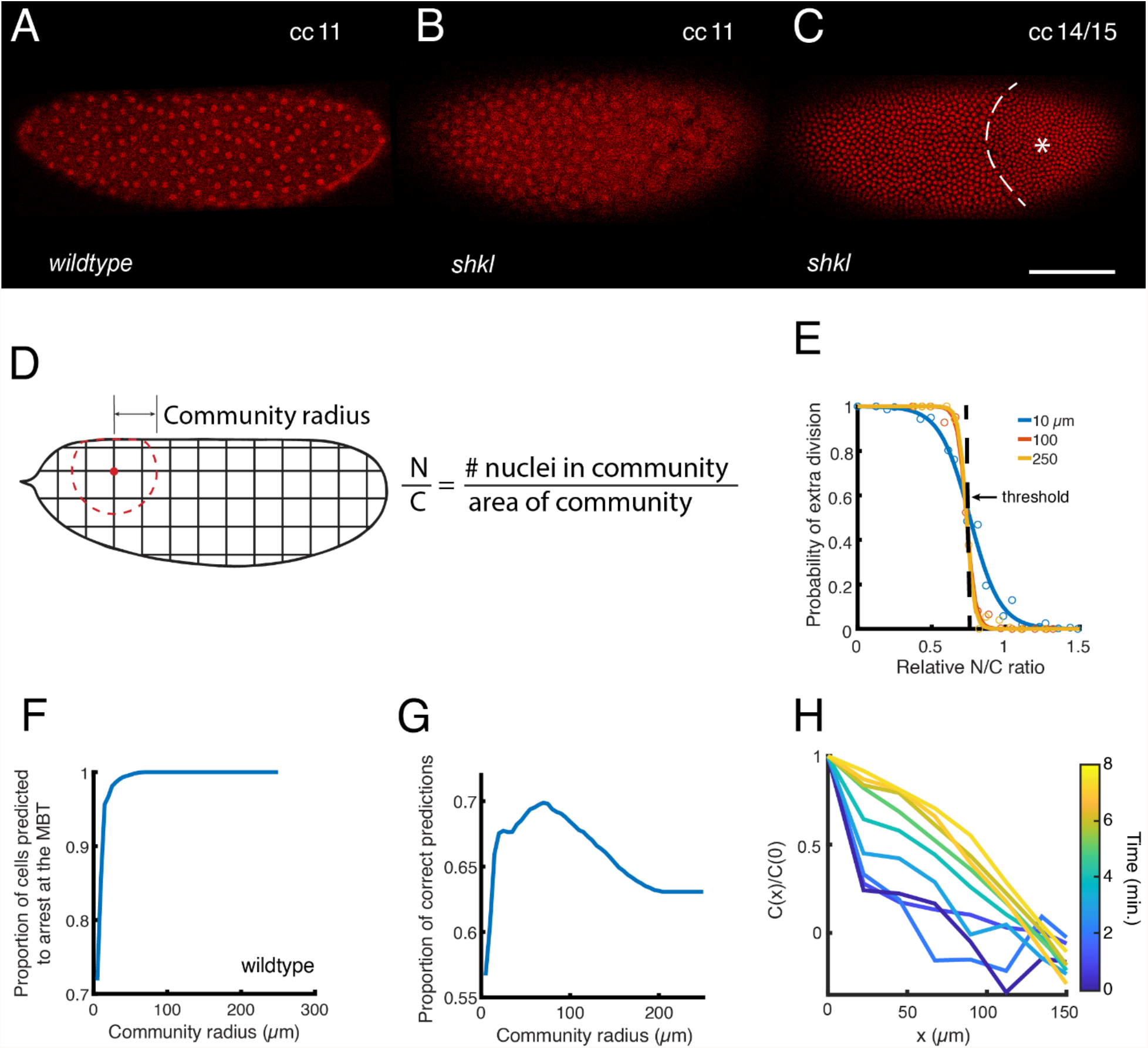
Sensing of the N/C ratio in *shkl* embryos. (A, B) Positioning of nuclei at cc 11 in wildtype (A) and *shkl* (B) embryos, visualized with His-RFP. (C) A *shkl* embryo at the MBT. The gradient of N/C ratio in *shkl* embryos frequently leads to the posterior undergoing an extra division. Dashed line: boundary between the normal and extra division regions. Asterisk: The region of the embryo which underwent an additional division. (D) Schematic of experimental design. Each embryo was discretized in a grid and a region with a certain community radius was specified. The N/C ratio was calculated as the number of nuclei in the community divided by the area. (E) The probability of a region dividing as a function of the local N/C ratio, relative to the average N/C ratio in wildtype embryos at cc 14. Curves for local neighborhoods of 10, 100, and 250μm are shown. A simple predictive model is a constant N/C ratio threshold, shown as a dashed line. (F) The proportion of nuclei in wildtype embryos which should arrest at the MBT as a function of the community radius. (G) Proportion of correct predictions in a test data set of *shkl* embryos using a simple threshold model as a function of community radius. (H) Two-point correlation function of the Cdk1/PP1 field as a function of distance for embryos in interphase of cell cycle 13. The correlation length was estimated as the point at which the correlation reaches 0.5 at the last time point and occurs at ∼100μm. Scale bar: 100μm.

**Fig. 4.**
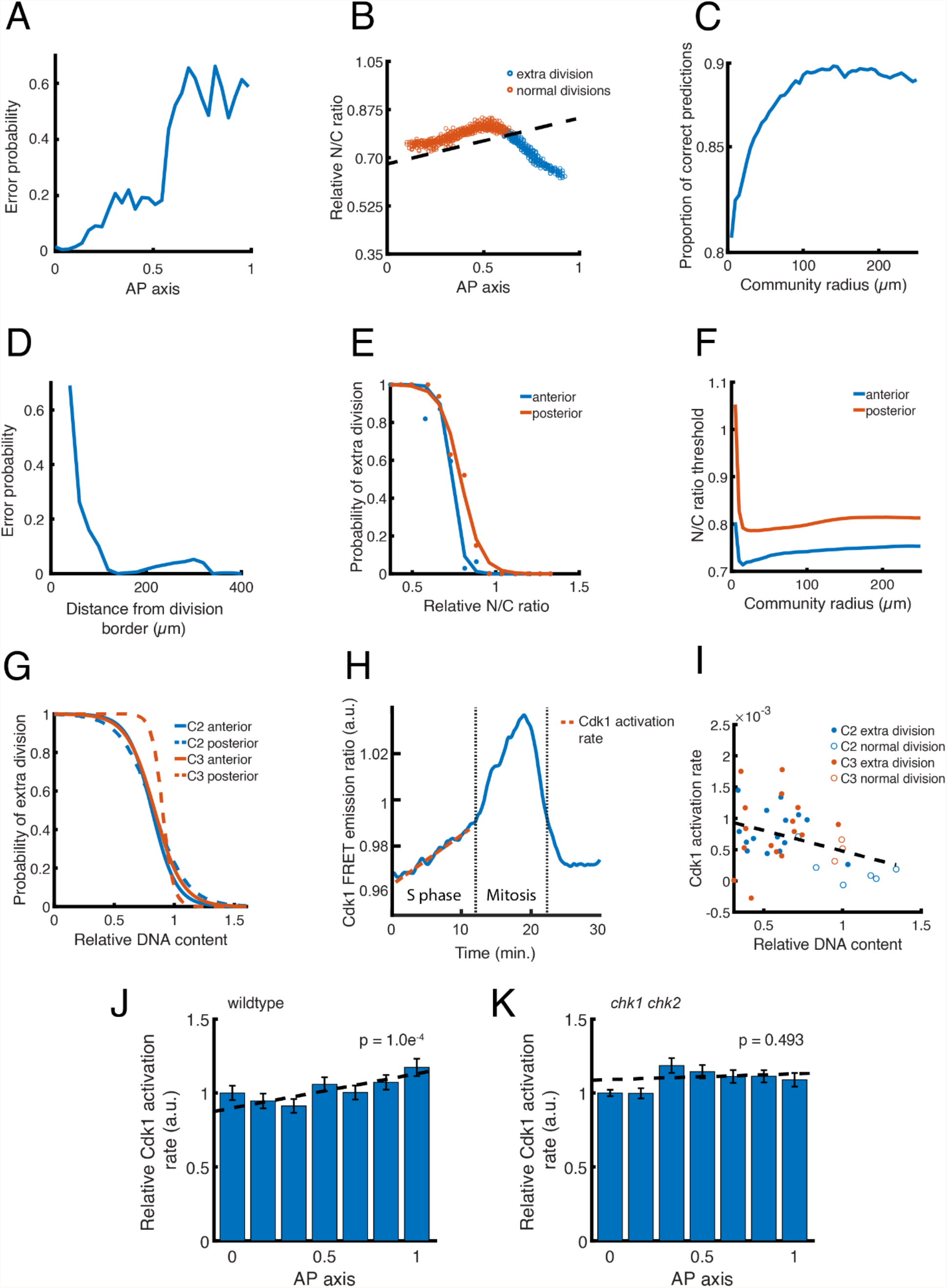
A gradient in N/C ratio sensing across the AP axis. (A) Errors made by the simple threshold model as a function of position across the AP axis. (B) The N/C ratio in local neighborhoods of 70μm (the maximum of the correct predictions of the simple threshold model) across the AP axis during cc 14. Dashed line shows that an N/C ratio threshold with a slight gradient across the AP axis does a much better job at dividing the normal and extra division regions. (C) This gradient threshold model predicts the division behavior correctly upwards of 90% of the time at a community radius of ∼100μm. (D) Errors in prediction from the gradient threshold model are largely near the border between normal and extra division regions. (E) Probability of a region undergoing an extra division as a function of N/C ratio in the anterior third versus posterior third of the embryo at a community radius of 100μm. The posterior curve is shifted towards a higher N/C ratio. (F) The N/C ratio threshold in the anterior and posterior of the embryo at different community radii. (G) Probability of a region undergoing an extra division as a function of N/C ratio at a community radius of 100μm, performed in embryos with either one additional or one fewer copy of either chromosome two or three. These embryos have altered DNA content but normal nuclear spreading. (H) Diagram showing an oscillation of Cdk1/PP1 FRET ratio during cc 13. The dashed red line indicates a linear fit of the slope of the ratio during S phase which we use as the Cdk1 activation rate. (I) Correlation between Cdk1 activation rate and division behavior in normal and extra division regions in compound chromosome embryos. At low DNA content and high Cdk1 activation rate, nuclei tend to divide. Dashed line: linear fit. (J, K) Cdk1 activation rate across the AP axis (relative to the anterior of the embryo) in wildtype (J) and *chk1 chk2* (K) mutants. Dashed lines indicate linear fit and p values (F-test) for the significance of the slope are shown. Data are represented as mean ± SEM.

Next, we tried to infer the optimal radius and a possible molecular mechanism for the collective nature of the cell cycle decision. Using the estimated thresholds, we measured the proportion of correct predictions made in a test data set of *shkl* embryos with regions of extra division and saw a peak in correct predictions at a community radius of 70μm (Fig. 3G). Next, we used the Cdk1/PP1 FRET biosensor to measure the correlation length of the Cdk1/PP1 activity field from the two-point correlation function. We found that this length (about 100 μm) is similar to the optimal community radius, thus suggesting that the collective decision of nuclei to undergo an extra division or not might reflect the fact that Cdk1/PP1 activity in neighboring nuclei influences each other (Fig. 3H). We have previously shown that spatial correlations in Cdk1/PP1 activity arises from the reaction-diffusion dynamics that drive the cell cycle during interphase.^24^ Thus, we conclude that the syncytial nature of the nuclear cycles coupled to the reaction-diffusion properties of the cell cycle oscillator ensure that nuclei act as large collectives and that such collective increase the robustness of the MBT.

### A gradient in sensing of the N/C ratio improves the ability to predict nuclear behaviors

While our previous results revealed the importance of collective N/C ratio sensing, we observed that the frequency of correct predictions from the model was limited to ∼70% (Fig. 3G), suggesting that we might be missing some additional regulation of the MBT or some additional aspects of the response to the N/C ratio. Thus, we sought to determine whether a more complex model, still centered on the N/C ratio, would account for the incorrect predictions (see Supplementary Figure 3). Notably, we found that the errors in prediction were distributed in a gradient across the AP axis (Fig. 4A), suggesting that nuclei might sense the N/C ratio differently depending on their location. We therefore hypothesized that a slight gradient in the N/C ratio threshold (with some small variance between embryos) would be a better predictor (Fig. 4B). With this model, we were able to correctly predict ∼90% of nuclear divisions with the best community radius of ∼100μm (Fig. 4C). Moreover, most of the errors in this model accumulated very close to the border of the normal and extra division regions (Fig. 4D), where we would expect that the ability of Cdk1 to propagate spatially might influence the decision.^10,27,28^

To further test the idea that the posterior of the embryo has a slightly higher N/C ratio threshold than the anterior, we compared the probability of nuclei dividing as a function of N/C ratio in the anterior and posterior. As predicted, the posterior third of the embryo showed a higher N/C ratio threshold than the anterior third (∼8% higher, Fig. 4E) which was independent of the community radius used (Fig. 4F). To control that the apparent gradient in N/C ratio response is not due to cytoskeletal and/or other effects of reduced Cul-5 activity, we investigated the decision of nuclei to arrest at the MBT when the N/C ratio in embryos (with uniform nuclear positioning) is brought closer to the threshold by genetic manipulations. We reasoned that in these embryos the proximity of the nuclear density to the threshold will result in a significant fraction of embryos having an extra division. To this end, we imaged embryos generated by crossing wild type females to males carrying compound chromosomes^25^. As a consequence, these embryos contained either one extra or one fewer copy of either chromosome 2 or 3. The embryos with one fewer copy of chromosome 2 or 3 have a DNA content (83% and 80% respectively) close to the 70% threshold seen in wildtype and also frequently had regions of an extra nuclear division.^25^ As in *shkl* embryos, the posterior of the embryos featured a slightly higher threshold than the anterior (∼2% increase for chromosome 2 and ∼8% increase for chromosome 3; Fig. 4G). In accordance, 100% (n=33) of extra divisions began in the posterior. Since the community radius and the correlation length of the Cdk1/PP1 field are both ∼100μm, we expect there to be a correlation between the Cdk1 activation rate measured during S phase (Fig. 4H) and nuclear density in the compound chromosome embryos. Indeed, we saw there was a correlation between the two, and the regions of the embryo which underwent an extra division were clustered at a low DNA content and higher Cdk1 activation rate (Fig. 4I).

To gain further insight on the slight gradient in the N/C ratio threshold across the AP axis, we divided wildtype embryos into grids and measured the Cdk1 activation rate in neighborhoods of each grid point at cc 13. We saw a slight but significant increase in the Cdk1 activation rate across the AP axis (Fig. 4J) which was not due to differences in the N/C ratio (Fig. S4). Since the increased duration of S phase at cc 13 is primarily due to the Cdk1 inhibition by Chk1^29-31^, we measured the Cdk1 activation rate in *chk1 chk2* mutants and saw that the gradient across the AP axis was ablated (Fig. 4K, S4). Thus, our results argue that the response of the cell cycle to nuclear density is not uniform across the embryo and that this difference might be dependent on the DNA replication checkpoint.

## Discussion

The tight control of the cell cycle and nuclear (cell) positioning and number is a ubiquitous feature of metazoan development and is crucial to the proper development of early embryos. A self-organized mechanism ensures uniform nuclear positioning across the AP axis in *Drosophila* embryos and thereby a synchronous halt at the MBT prior to gastrulation.^3^ In this work we have taken advantage of *shkl* mutants which have defects in nuclear spreading to identify a novel pathway involved in the control of cortical contractility and gain insights into how nuclei respond to changes in the N/C ratio.

Through DNA sequencing and complementation tests, we have identified *shkl* mutants as mutations of the ubiquitin ligase Cul-5. In the early embryo, Cul-5 does not regulate the cell cycle oscillator but is required for Rho and myosin activity. In many systems Cul-5 restricts the levels of active Src kinase^13-15^, which is a known regulator of the actomyosin cytoskeleton. Indeed, we found that the *cullin-5* phenotype could be largely rescued through a genetic reduction in Src activity and recapitulated through Src overexpression, indicating that a main function of Cul-5 is to downregulate Src activity. These results implicate the Cul-5/Src axis as a crucial pathway involved in the control of cortical contractility in early *Drosophila* embryos.

In the early embryo, nuclei regulate their own positioning through PP1 activity which spreads from the nuclei to the cortex.^3^ This localized PP1 activity drives activation of Rho and Myosin II accumulation in turn.^3^ Our results argue that Cul-5 and Src act in a pathway downstream or parallel to the cell cycle to regulate Rho activity. The molecular mechanisms by which Cul-5 and Src control Rho remain to be elucidated, as is the possible connection between the cell cycle oscillator and Cul-5/Src activities. Since Src has been shown to regulate Rho GTPases in several contexts via the control of guanine nucleotide exchange factors and/or GTPase-activating proteins,^32-34^ these proteins are natural candidates for the regulation of cortical actomyosin regulation via the Cul-5/Src pathway.

Control of the MBT by the N/C ratio is important in several species, including *Drosophila* and *Xenopus*^25,26,35-38^ but likely excluding zebrafish.^39-42^ This density of DNA (as well as nuclear size^43,44^) can directly or indirectly impact multiple aspects of the MBT, namely zygotic gene expression^45-48^ and cell cycle control.^26,35,49-51^ Here, we have exploited the changes in nuclear positioning in *shkl* embryos to generate a continuous range of nuclear densities. This property allowed us to gain insights into how the decision of nuclei to pause their cell cycles at the MBT is affected by the N/C ratio. We found that the threshold for nuclear division is about 70% of the density at nuclear cycle 14, which confirms previous results. This value—halfway between the density at cycle 13 and 14—likely contributes to the robustness of the MBT. However, this value is not sufficient for the robustness of the MBT. To ensure reliable lengthening of cycle 14 in all nuclei, the sensing of the N/C ratio must be averaged over hundreds of nuclei. Consistently, our results indicate that nuclei might sense the local N/C ratio in neighborhoods of ∼100μm. This length essentially coincides with the correlation length of the Cdk1 activity field, which is established via reaction-diffusion mechanisms.^24^ Additionally, we found that a model based on uniform sensing of the N/C ratio fails to predict the behavior of a large fraction of nuclei (only 70% of nuclei are predicted correctly). However, a model assuming a slightly higher N/C ratio threshold in the posterior is highly predictable (>90% prediction ability) and mainly misses the behavior of nuclei at the interface between the region of extra division and that of normal division. Thus, we propose that the N/C ratio is the major regulator of the cell cycle at the MBT and that no mechanism other than a slight spatial modulation of the N/C threshold is needed to account for nuclear behaviors. This spatial modulation likely reflects the fact that the rate of Cdk1 activation is also slightly graded across the AP axis. The Cdk1 activation gradient is dependent on the DNA replication checkpoint, which argues that the gradient might be controlled by an asymmetric distribution of factors controlling DNA replication and/or Chk1 activity.^52-54^ Alternatively, the DNA replication checkpoint and Cdk1 activity might be influenced by factors controlling AP patterning and expressed in gradients across the embryos.^55^ In the future, it will be interesting to understand the mechanisms and possible functional significance of this gradient.

The precise coordination of biochemical and mechanical signals is a ubiquitous feature of embryonic development. In early *Drosophila* embryogenesis, it is necessary for the uniform positioning of nuclei and timing of the MBT. Our work has identified a new pathway wherein Cul-5 regulates cortical contractility by restricting Src activity. Our results investigating embryos with patchy divisions indicate that nuclei sense the N/C ratio in neighborhoods of ∼100μm and pause the cell cycle when the local density exceeds a threshold around 70% of the normal density at the MBT. Moreover, the threshold required to arrest the cell cycle is slightly graded across the AP axis and is coupled to the spatiotemporal dynamics of Cdk1. Quantitatively measuring biochemical and physical dynamics during specific morphogenic events will undoubtedly continue to reveal new insights into the mechanisms and regulations of these pathways.

## Acknowledgments

We thank the Bloomington Drosophila Stock Center, the Kyoto Drosophila Stock Center, Ruth Lehmann, Denise Montell and Alana O’Reilly for providing stocks. We thank the Drosophila Genomics Resource Center for constructs. We acknowledge discussions with Massimo Vergassola. We thank members of the Di Talia lab for comments on the manuscript. This work was supported by a Schlumberger Faculty for the Future Fellowship and an HHMI International Student Research Fellowship to V.D., Associazione Italiana Ricerca sul Cancro (AIRC) MFAG-2020 n. 25040 to A.P.; University of Torino Fondo Ricerca Locale 2019 and 2020 PULA_RILO_19_01 and PULA_RILO_2020 to A.P., and NIH (R01-GM122936 and R01-GM136763) to S.D.

## Author contributions

Conceptualization, L.H., A.C., V.E.D., and S.D.; Methodology, L.H., A.C., V.E.D., A.P., and S.D.; Software, L.H. and A.P.; Investigation, L.H., A.C., V.E.D., and A.P.; Writing – Original Draft, L.H. and S.D.; Supervision, A.C and S.D. Funding Acquisition, S.D.

## Supplementary Figures

**Fig. S1.**
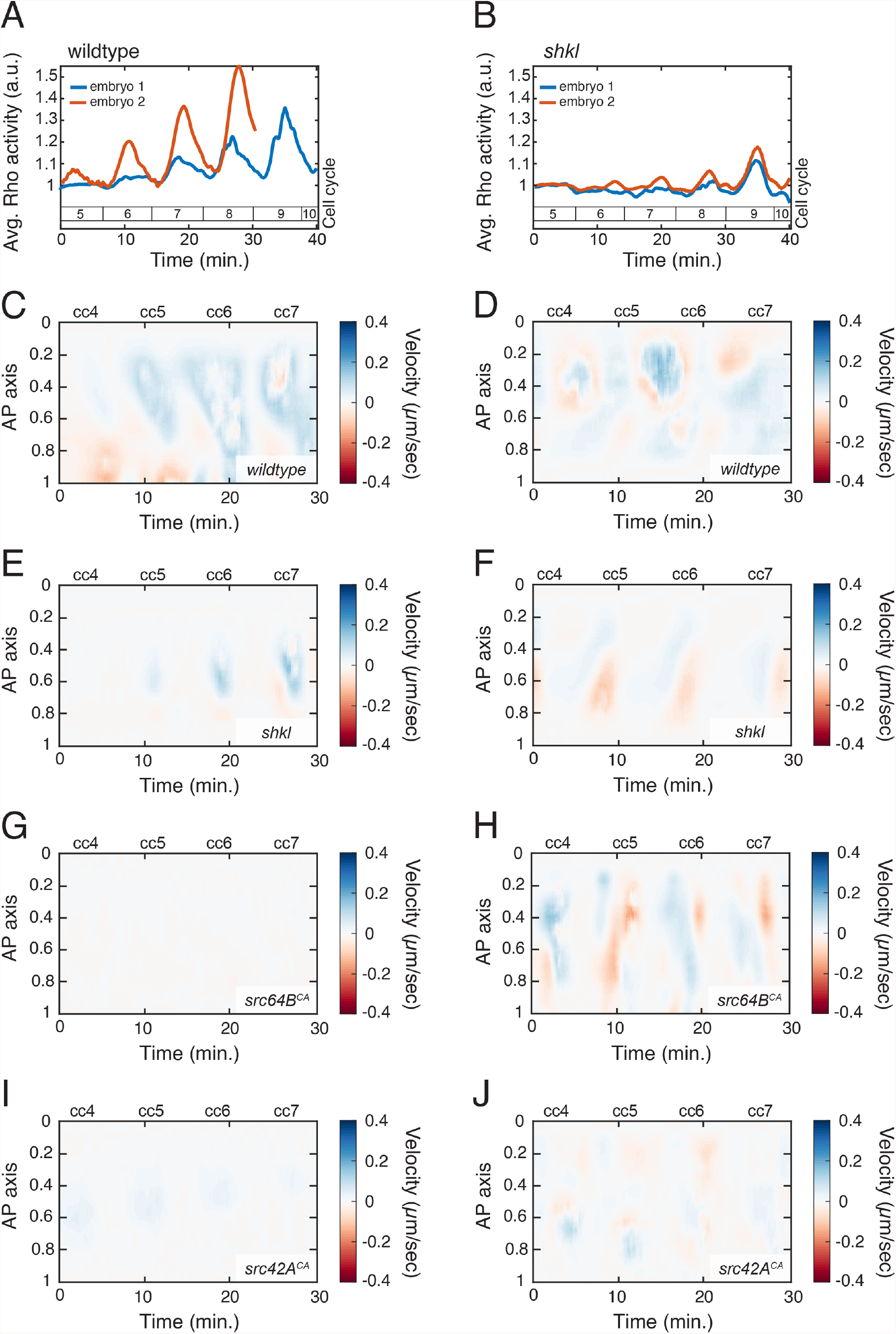
Quantification of cortical contractility and cytoplasmic flows. (A, B) Replicate quantification of Rho activity in wildtype (A) and *shkl* (B) embryos. (C-J) Replicate heatmap quantification of cytoplasmic flows in wildtype (C, D) *shkl* (E, F), src64B overexpression (G, H), and src42A overexpression (I, J).

**Fig. S2.**
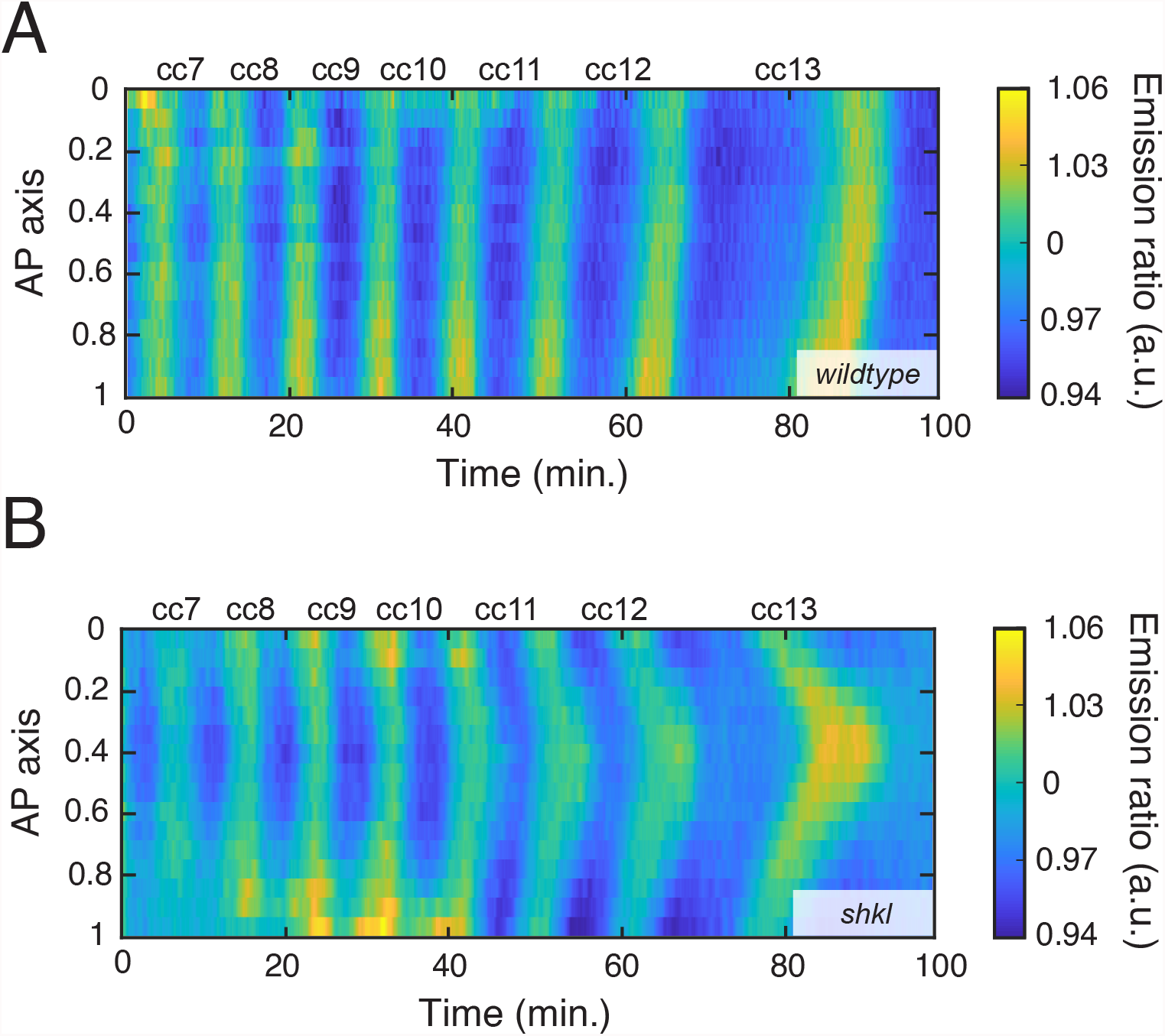
Spatial quantification of Cdk1/PP1 FRET sensor. Cdk1/PP1 FRET emission ratio for wildtype (A) and *shkl* (B) embryos, measured across the AP axis and time. In time, oscillations are unperturbed, and cell cycle duration is unchanged. The lower nuclear density in the posterior relative to the anterior lead to slightly earlier divisions and a gradual asynchrony in cell division builds up over time.

**Fig. S3.**
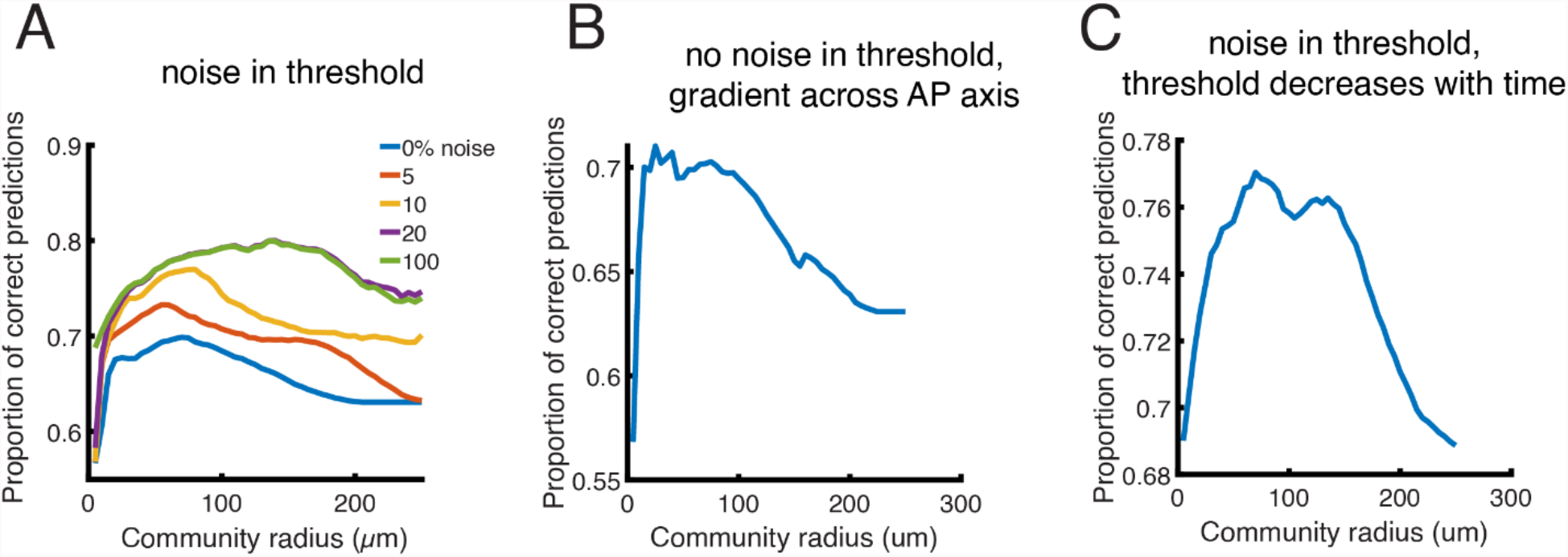
Additional models of N/C ratio sensing. (A-C) Proportion of correct predictions in possible models of N/C ratio sensing. (A) The decision to pause the cell cycle is determined by a global threshold of ∼70% (see main text Fig. 3E) which varies up to a certain percentage of this threshold (colored lines). Allowing “noise” of 10% means the threshold for that embryo must fall in the range [1/1.1 to 1.1]. (B) The decision to pause the cell cycle is determined by a global threshold (Fig. 3E) with an additional gradient in threshold of up to 10% across the AP axis. (C) The decision to pause the cell cycle is determined by a global threshold with noise (as in panel B). Additionally, when the first nucleus divides, the threshold lowers at some rate over time (tested uniformly from 0 to up to 15%/minute), making it more difficult for nuclei to divide.

**Fig. S4.**
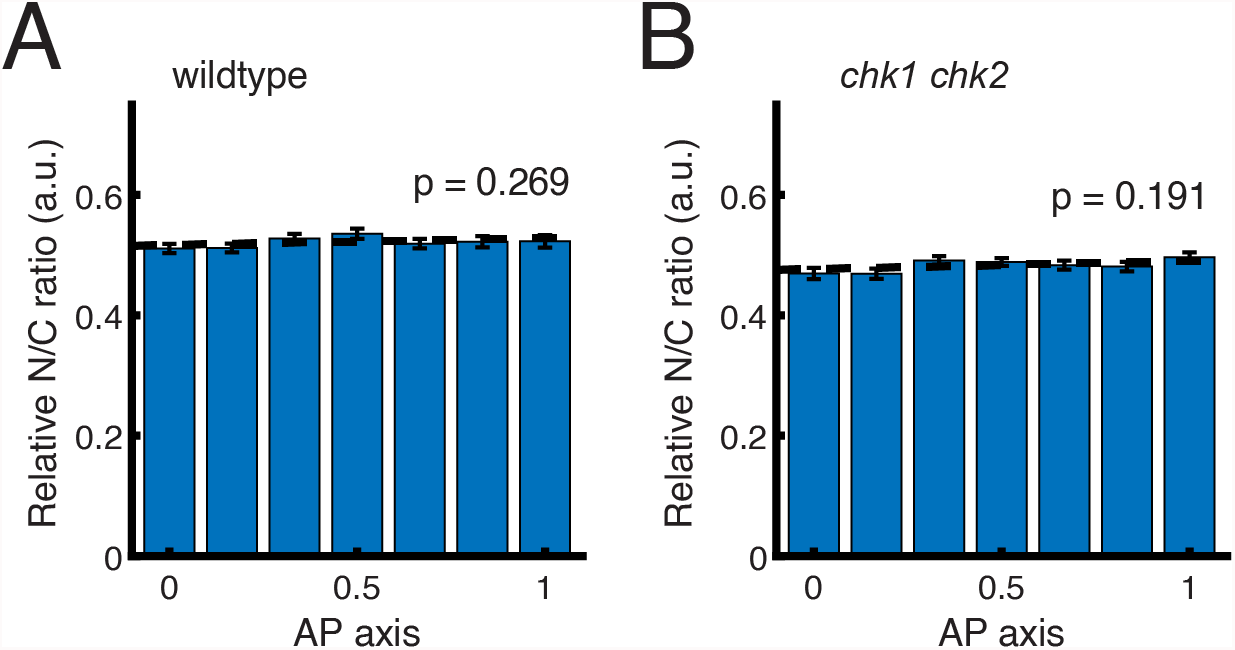
Nuclear density across the AP axis. (A, B) N/C ratio (relative to the embryo’s anterior) across the AP axis in wildtype (A) and *chk1 chk2* (B) embryos. P-values (F-test) for the significance of the slope are shown and indicate a uniform nuclear density across the AP axis for both genotypes.

